# Satiety does not abolish Pavlovian-to-instrumental transfer but accelerates devaluation in humans

**DOI:** 10.1101/2025.07.22.666116

**Authors:** Marc Ballestero-Arnau, Borja Rodríguez-Herreros, Manuel Moreno-Sánchez, Antoni Rodríguez-Fornells, Oliwia Trembecka, Toni Cunillera

## Abstract

Environmental cues that predict palatable food can robustly drive behavior, override satiety signals and promote maladaptive eating habits. However, conventional human paradigms often fail to capture the dynamic interplay between hunger, satiety, and cue-induced behavioral actions. To address this, we built a novel automated dispenser synchronized with task events and integrated into an adapted human Pavlovian-to-instrumental transfer (PIT) paradigm to deliver real, consumable food rewards in real time—allowing participants to eat online and thereby induce satiation or sustain hunger. In one experiment, we manipulated food rewards by varying portion size and consumption timing. Large portions consumed immediately induced rapid satiation, whereas small, trial-by-trial portions kept participants non–satiated; correspondingly, large portions accelerated devaluation of the food reward—evidenced by a pronounced decline in instrumental responding—while small portions sustained cue-driven behavior over time. In a subsequent experiment, small, immediately consumed food rewards (which kept participants non–satiated) were directly compared with real-time monetary rewards. Although both reward types initially invigorated behavior, the motivational impact of monetary rewards declined quicker, suggesting that the absence of physiological feedback renders secondary reinforcers more vulnerable to rapid devaluation. These findings suggest that satiety accelerates, rather than abolishes, the influence of food-associated cues. The distinct temporal dynamics observed for food and monetary outcomes underscore the importance of real-time ingestion in human PIT paradigms, offering new insights into the mechanisms by which physiological states modulate cue-driven devaluation.

## Introduction

In daily life, we are frequently exposed to environmental cues that predict palatable, high-calorie foods, ranging from the aroma of pastries to the logos of sweet drinks or processed snacks (Watson et al., 2014). In environments where palatable foods are constantly available, such cues can override internal satiety signals, promoting excessive intake and contributing to maladaptive eating patterns such as obesity (Swinburn et al., 2011; van den Akker et al., 2018). Interestingly, maladaptive food-seeking behaviors have been suggested to mimic addictive patterns, as Pavlovian cues can sustain motivational responses even when the reward value is diminished (Garbusow et al., 2022; Havermans, 2013; Robinson et al., 2015). This persistence is evident, for example, when food-paired cues continue to enhance instrumental responding despite outcome devaluation due to satiation (Colagiuri & Lovibond, 2015; De Tommaso et al., 2018). For example, visually salient cues such as colored packaging, food advertisements, or the strategic placement of snacks may trigger approach behaviors, prompting individuals to seek and consume food even when they are no longer hungry. Understanding how these cues drive approach behaviors despite devaluation is crucial for developing interventions aimed at reducing maladaptive eating patterns.

Building upon this phenomenon, early foundational work by Estes (1943) in animals revealed that a stimulus that predicts food can invigorate reward-seeking behaviors (e.g., lever-pressing), illustrating how Pavlovian associations enhance instrumental responding (Marshall et al., 2018). Many studies have consistently replicated these observations in both animal and human settings (Burghoorn et al., 2024; Corbit & Balleine, 2005; Holland, 2004; Rescorla, 1994; Talmi et al., 2008). Indeed, recent meta-analyses and reviews (Cartoni et al., 2016; Garbusow et al., 2022; Holmes et al., 2010) suggested that these cue-driven effects are remarkably robust and extend across species, emphasizing the potency of Pavlovian cues in modulating instrumental behavior.

Importantly, classic rodent paradigms typically involve delivering real food rewards in small, repeated portions throughout each session, allowing hunger and satiety to fluctuate. Under these conditions, animals naturally undergo real-time devaluation of the reward as they become sated (Corbit et al., 2007; Holland, 2004). This animal model design captures a crucial feedback loop: as hunger decreases and satiety accumulates, reward-seeking behavior decreases on a within-session basis (Corbit & Balleine, 2005). In these models, the interplay between reward ingestion and emerging satiety precisely explains when and why cues lose their capacity to invigorate reward seeking (Balleine & Dickinson, 1998; Corbit & Balleine, 2005; Holland, 2004). However, many human experiments diverge from this approach by substituting tangible foods with images, tokens, or hypothetical rewards (Cartoni et al., 2016; Eder & Dignath, 2016; Hogarth & Chase, 2011; Talmi et al., 2008). Although these substitutes offer practical advantages—such as easing experimental procedures and avoiding dietary restrictions (Allman et al., 2010; Eder & Dignath, 2016; Hogarth & Chase, 2011)—they also eliminate the actual consumption that could trigger real-time satiety signals (Pool et al., 2016). Consequently, participants may maintain artificially elevated motivation or craving because they never experience the gradual shift from hunger to satiety that characterizes rodent experiments (Marshall & Ostlund, 2021). This methodological gap can generate mixed or attenuated findings, as real-time physiological feedback – integral to rodent studies– remains largely absent.

Although a handful of human Pavlovian-to-instrumental transfer (PIT) studies have examined devaluation—most notably Colagiuri and Lovibond (2015), who used pre–feeding to reduce transfer—none have delivered the actual reward online during testing. To fill this gap, we built an Arduino–controlled dispenser that drops real snack portions in sync with task events, allowing us to vary portion size (large vs. small) and consumption timing (immediate vs. delayed) on a trial–by–trial basis. This setup let us induce rapid satiation or maintain non–satiated states and then directly measure how these physiological states influenced cue–driven responding. Such real–time ingestion aligns human protocols more closely with animal work (Balleine & O’Doherty, 2010; Cartoni et al., 2016; Pool et al., 2016). For example, Pool et al. (2016) proposed that introducing actual food—as opposed to pictures—triggered a more pronounced shift in attention and approach behaviors once satiation began, underscoring the importance of embodied physiological signals (e.g., gastric distension, hormonal feedback).

With this device in place, we implemented two animal–derived manipulations. First, consumption timing: rodents eat immediately, so within–session satiety rapidly devalues later rewards (Corbit et al., 2007; Holland, 2004), whereas delaying human consumption may artificially sustain motivation (Pool et al., 2016). We therefore compared trial–by–trial versus delayed consumption to assess how immediate versus delayed ingestion shapes learning and PIT transfer. Second, portion size: larger quantities accelerate fullness and should accelerate devaluation (Lingawi et al., 2022), since even modest satiety gains blunt approach in animals (Panayi & Killcross, 2022). Therefore, by testing small versus large portions in humans, we examined how quickly reward–seeking declines as physiological satiety accumulates.

As mentioned, one way to explore whether reward-seeking behaviors are downregulated is by investigating cue–outcome devaluation—the decline in responding that occurs when an outcome loses its subjective value (e.g., due to satiety or outcome availability). In the present study, we employ “devaluation” to refer to the behavioral decrease that emerges when a Pavlovian cue no longer drives responding because the food has become less appealing (e.g., through satiety), whereas “extinction” is used to describe our specific method of removing rewards during certain phases (transfer) of the study (Seabrooke et al., 2018). To this end, the PIT paradigm provides a well-established framework to disentangle how Pavlovian cues modulate independently learned instrumental actions (Corbit et al., 2007; Holland, 2004). In a typical PIT design, participants first undergo Pavlovian training, in which a neutral cue is paired with a food reward. Separately, they learn an instrumental response—such as pressing a key— to earn the same or a different food outcome. During the critical transfer test, the previously learned Pavlovian cue is presented while the participants must perform the instrumental action again. Differences in response rates (number of keys pressed) when the Pavlovian cue is present provide evidence that the cue invigorates instrumental behavior (Lingawi et al., 2022; Talmi et al., 2008). In the present PIT paradigm, participants used repeated key responses (during a 3 seconds window) that were consistently paired with a single food reward, which aligns with general PIT models (Lingawi et al., 2022). This design mirrors animal paradigms in which a single lever press is paired with one outcome, producing general motivational effects (De Tommaso et al., 2018). As Cartoni et al. (2016) emphasized, single-reward designs are well suited for examining how outcome devaluation—such as that induced by satiety—modulates general incentive motivation. Furthermore, other findings suggest that general PIT is particularly sensitive to devaluation through satiety, supporting the relevance of the general PIT design for investigating how satiety influences cue-driven behaviors (Lingawi et al., 2022).

Overall, given evidence that the PIT paradigm can elicit devaluation effects in humans, a deeper understanding of how satiety contributes to the devaluation of the same food item is still necessary. Accordingly, the present study investigates whether real-time satiety—induced by immediate or delayed consumption of small or large food portions—attenuates or otherwise alters PIT in humans and whether satiety accelerates the devaluation of food rewards (Panayi & Killcross, 2022).

## Experiment 1

To examine how reward magnitude (small vs. large portions—designed to maintain non–satiation or induce rapid satiety) and consumption timing (immediate vs. delayed) interact, we employed an adapted appetitive PIT paradigma (da Costa et al., 2020; Estes, 1943; Marshall & Ostlund, 2021). In the Pavlovian phase, participants learned cue–food associations, followed by an Instrumental Go-No-Go phase where they earned the same food rewards and finally a transfer phase in which the Pavlovian cues were reintroduced in an extinction context (Meemken & Horstmann, 2019).

To this end, we implemented a between-subjects design with four conditions, created by orthogonally crossing portion size (small vs. large) with consumption timing (immediate vs. delayed). In the small reward–immediate (SRI) condition, participants received small, consumable portions on a trial-by-trial basis, thereby slowing the buildup of satiety. In contrast, the large reward–immediate (LRI) condition involves the immediate consumption of larger portions, which we expect to induce a faster onset of satiety. In the SRD and LRD conditions, rewards accumulated until the end of the block, therefore postponing consumption—and satiety—until the transfer phase of the PIT paradigm. We hypothesized that during the learning phases (Pavlovian and Instrumental), all groups would generate robust cue–food and response– food associations. However, during the transfer test, participants in the LRI condition are expected to show the most pronounced decline in cue-elicited responding due to rapid satiety, whereas those in the SRI group should continue responding for a longer period. Under delayed consumption conditions (SRD and LRD), we anticipate that postponement may either preserve motivation by delaying satiety-related devaluation or reduce subjective reward value through temporal discounting. Because delay discounting is supposed to be more pronounced for smaller rewards (Odum et al., 2006), we expected the SRD group to show somewhat faster devaluation under postponement. This design allows us to clarify how real-time satiety, portion size, and consumption delay converge to shape the persistence of food-seeking behavior as a function of physiological (hunger/satiety) modulation.

### Methods

#### Participants

Ninety-nine students (83 female) from the Faculty of Psychology at the University of Barcelona participated in the experiment [mean age = 21.56 years, S.D. = 3.2, range = 18–42]. A power analysis using G*Power (Erdfelder et al., 2009) indicated that a sample of 88 participants was sufficient to assess the effect of 2 within-subjects variables and 2 between-subjects variables (with a power = 95% and an a priori alpha set at p = 0.05) for a moderate effect size estimate (η*_p_*^2^ = 0.06). All participants had BMIs within the normal range (18–25) [mean BMI: LRI = 21.24, S.D. = 1.8; LRD = 21.09, S.D. = 2.3; SRI = 21.48, S.D. = 1.9; SRD = 21.15, S.D. = 1.9; *F*(3,95) = 0.17; *p* = 0.917; η*_p_*^2^ = 0.05]. Prior to the sessions, all participants completed at least 16-hour fasting period (reported by the participants at the beginning of the session) to maintain high motivational salience of food (Finlayson et al., 2007). These fasting hours were similar between conditions [LRI = 16.88, S.D. = 1.4; LRD = 17.11, S.D. = 1.9; SRI = 16.66, S.D. = 0.9; SRD = 16.47, S.D. = 0.9; *F*(3,95) = 1.01; *p* = 0.391; η*_p_*^2^ = 0.03]. All participants provided informed consent as approved by the local ethics committee (IRB00003099) before their participation and received 20€ for completing the experiment.

#### Materials & Procedure

##### Preliminary measurements

We measured blood glucose concentrations in all participants twice, first upon their arrival at the laboratory and then after completing the experiment, using a glucometer (SD Biosensor Inc.) following a standardized safety protocol. The participants were subsequently asked to evaluate their subjective measures of hunger using a visual analog scale (VAS) of 10 cm, ranging gradually from 0–10, where 0 meant *not hungry at all* and 10 corresponded to *very starving*. We then assessed participants’ liking of the food reward used in the experiment while informing them about the number of reward units they could gain and eat during the experiment. They were then asked to taste a sample of a food unit (extruded cornmeal snack) and respond to the question *“How much do you like this cornmeal snack as a reward during the experiment*?” using an analogous VAS (where 0 meant *totally dislike* and 10 *like the most*). Finally, we required all the participants to report their immediate subjective desire (wanting/craving) to receive and eat (cornmeal snacks) the reward item of the experiment, again using a 10 cm VAS (0 indicated *no desire at all* and 10 *extreme desire*).

##### Reward delivery machine

To ensure synchronisation between the delivery of the reward and the visual feedback presented on the screen during the corresponding rewarded trials throughout the experiment, we developed a device operated by an Arduino and programmed with PsychoPy (2022.2.4 version; Peirce et al., 2019). In all rewarding trials, the device moved a tray horizontally to dispense as many reward units as needed. The machine was equipped with a tray grid containing 100 cube-shaped slots, each measuring 1 cm^3^, designed to hold 100 individual reward units. The grid was replenished during pause between blocks in both the Pavlovian and Instrumental tasks. The grid moves horizontally along either the x- or y-axis across a 55 *×* 40 cm platform and is positioned 38 cm above the table surface. The platform included a circular opening with a diameter of 23 mm in its center through which the reward units were dispensed. After dispensing, the reward units travelled through a PVC tube to a plate on the table, located approximately 62 cm to the right of the participants’ right arm. A reward unit reached the plate approximately 700 ms after the onset of the corresponding visual feedback and was accompanied by an intrinsic sound of approximately 53 db.

##### Pavlovian task

The Pavlovian task comprised the presentation of a fractal image followed by a musical tone in each trial, with only one out of four combinations (25% of the trials) being associated with the possible appearance of a reward. All four visual–auditory combinations consisted of either blue or orange hue square shape fractal pattern images of 90 × 90 mm, whereas the tones corresponded to the G musical tone and the raised semitone variation of this tone G♯ (G Sharp), both played by string instruments corresponding to a classical guitar and a ukulele. The four tones were generated using a Python script and a specific package for tone generation [‘pydub’; Version 0.23.1] and had a duration of 300 ms with 50 ms ramps on and off. The two fractal images were each assigned to a single instrument and paired with two musical tones (G and G♯) played by that instrument. Image 1, for example, was always accompanied by the guitar playing G or G♯, while Image 2 was always accompanied by the ukulele playing G or G♯. This arrangement yielded four unique image–tone compounds, of which one was randomly designated as the rewarded conditioned stimulus (CS+) and the other three served as nonrewarded stimuli (CS–). Using two similar pitches on two different instruments created a challenging discrimination task and kept participants engaged during passive Pavlovian learning. In each trial, feedback was always provided after the presentation of the CS compound. Positive feedback was presented in the CS+ rewarded trials (winner trials) and negative feedback was delivered in the CS- trials and not rewarded in the CS+ trials.

The reward consisted of either one (small-reward conditions) or two (large-reward conditions) food items, delivered in synchronization with the presentation of positive feedback on the screen and consumed during the task. A reward was provided in 70% of the trials where the CS+ was presented (see Figure 1A and the Reward Delivery Machine section). A probabilistic reward delivery procedure was employed to enhance participant engagement in the task (Holland, 2004). Participants were instructed to remain attentive throughout the task to identify the rewarded audio-visual compound—the CS+—as they would be tested on this learning at the conclusion of the task.

**Figure 1:**
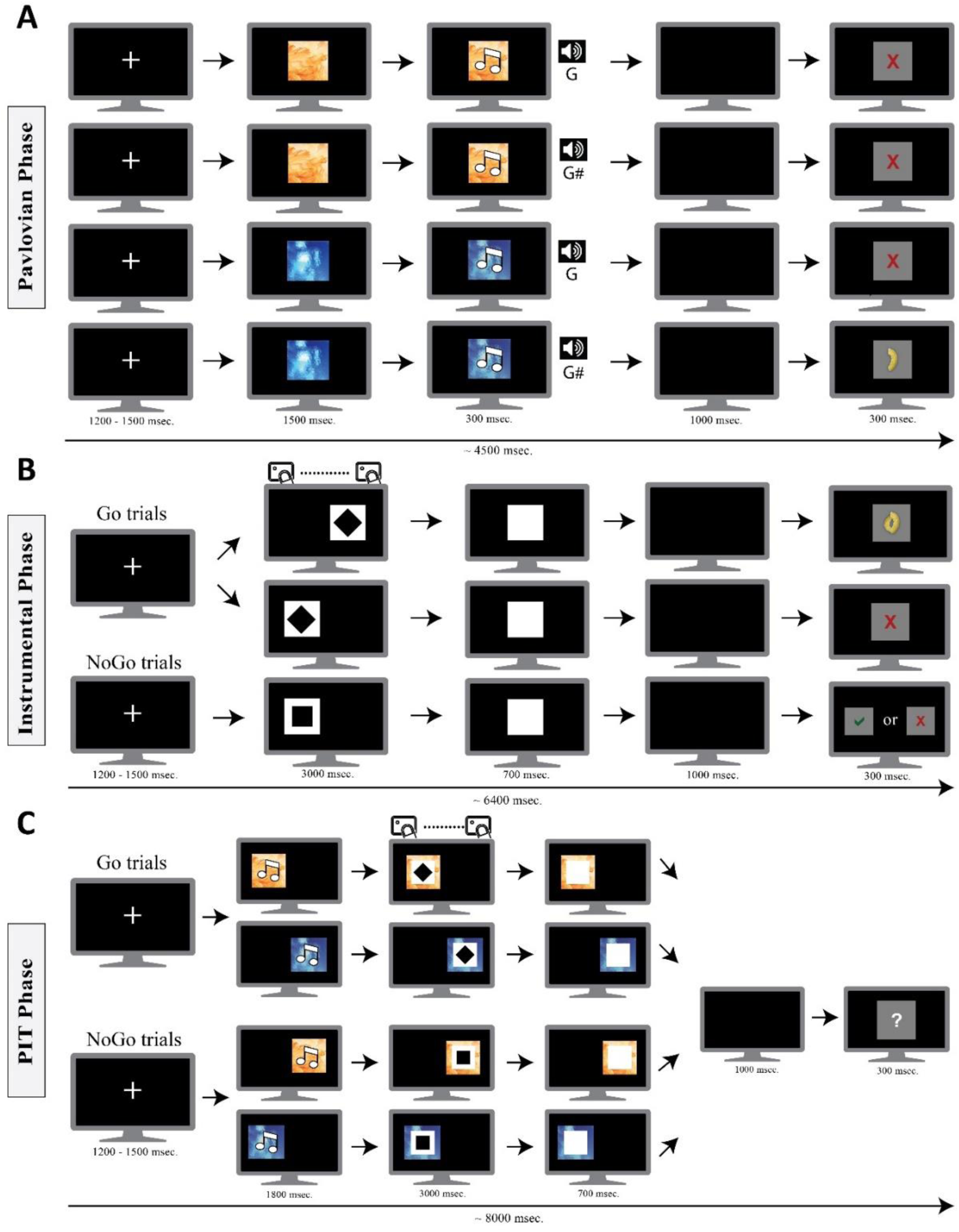
**A**) Flow representation from the Pavlovian task: Each trial began with a jittered fixation cross (between 1200–1500 ms), followed by a fractal image. A tone was played 1500 ms after image onset for 300 ms. A black screen (1000 ms) followed before positive or negative feedback was presented, along with the reward units (food in experiment 1; food or Money in experiment 2) in the case of a winner trial (300 msec). **B**) Instrumental task: Each trial began with a fixation cross (jittered between 1200–1500 msec). Subsequently, a Go-NoGo signal appeared (left or right side of screen), superimposed on a white square for 3000 ms, where participants responded or withheld responses depending on the Go-NoGo signal. After that, the white square remained for an additional 700 ms poststimulus. A black screen (1000 msec) followed, and then positive or negative feedback was presented, along with the reward units (food in experiment 1; food or Money in experiment 2) in the case of a winner trial (300 msec). **C**) Flow of the transfer task, where the two previous tasks were combined and the Go-NoGo signals were presented superimposed on the Pavlovian image after the tone presentation. The only difference was the uninformative feedback used during the whole task.

The Pavlovian task consisted of 6 blocks of 40 trials (10 trials for each CS compound). The first 5 blocks (200 trials) were designed as a passive learning task in which participants were instructed to observe and discover the image–sound combination that was associated with the delivery of the reward, whereas the last block was designed to obtain indirect measurements of CS+ learning. In all six blocks, each trial began with a fixation cross displayed for a jittered duration between 1200 and 1500 ms. Subsequently, a fractal image was presented for 1800 ms. After 1500 ms of the onset of the image presentation, a tone was played for 300 ms, with both the image and tone ending 1800 ms after the image onset presentation. Immediately after, a black screen was displayed for 1000 ms, followed by the presentation of the feedback for 300 ms, which was synchronized with reward delivery in rewarded trials (see Figure 1A). Participants received feedback in all trials consisting of the amount of reward items obtained in each trial. A total of 7 (small-reward condition) or 14 (large-reward condition) reward items were delivered in each block (a total of 42 small reward or 84 large reward units for the whole task). All the stimuli in the Pavlovian task were presented on a screen subtending approximately 10.4 degrees of the visual angle.

In the sixth block of the Pavlovian task, the feedback displayed on the screen was covered by a patch. The participants were informed that they could either press a pedal with their foot to immediately reveal the feedback or wait 3500 ms for the feedback to be automatically displayed. Before the block commenced, the participants were asked to remove the shoe from their dominant foot to enhance their tactile sensibility to the response device. They were also instructed on the new specificities of the task. This new procedure allowed us to record both the latency of pedal presses and the ratio of responses. Compared with those of the CS-, lower latencies for the CS+ and higher pressing ratios were considered indicative of learning the reward contingency (Bucker & Theeuwes, 2017). Consistent with animal PIT studies, we took special precautions to differentiate the motor actions required for the Pavlovian and Instrumental tasks (Corbit & Balleine, 2003). Whereas the Pavlovian task involved pressing a pedal with one foot in the last block, the Instrumental task required continuous keyboard presses using the index fingers of both hands. This ensured that motor preparation for the Pavlovian and Instrumental responses remained distinct, thereby reducing potential confounding factors.

##### Explicit Pavlovian learning test

After completing the Pavlovian task, the participants performed a test consisting of eight trials. During this test, each CS audio-visual compound was presented twice. The participants were instructed to use the left mouse button to indicate whether each compound was a rewarded or a nonrewarded one. Each trial followed the same structure as in the Pavlovian task, except that feedback was not presented. Instead, two images of buttons labeled “Yes” and “No” appeared on the left and right sides of the screen, respectively. The participants were instructed to click “Yes” if the compound had been previously associated with a reward and “No” if it had not. The Yes/No button images remained on the screen until a response was made. No feedback was provided during the test.

##### Instrumental task

The next task of the experiment consisted of an Instrumental learning task combined with a Go-NoGo task (Freeman et al., 2014). In each trial, one of two different geometric figures was presented on either the left or right side of the screen. The two figures consisted of the same black square (30 × 30 mm), which could be displayed either nonrotated or rotated 45°, making it appear as a rhombus (see Figure 1B). The figures were always superimposed on a larger (60 × 60 mm) white square. Before starting the task, each participant was randomly assigned one figure as the Go signal and the other figure as the NoGo signal. The participants were instructed that correct responses to the Go stimulus would be rewarded. The Go signal was presented on 75% of the trials, while the NoGo signal was presented on 25% of the trials. The presentation order of those signals was randomized for each block, with the only constraint being that a NoGo trial could not be presented more than 3 times in a row. The correct response pattern required to obtain the reward was defined before commencing the experiment, using a specific training protocol. The output of this training provided the average number of responses made in front of the Go signal, which was then fixed as the individual’s baseline response pattern.

The training consisted of a short and simplified version of the Instrumental task, in which for every trial, the same Go signal (a black triangle) was presented over a white 60 × 60 mm square. Participants were required to correctly perform 10 out of 16 practice trials to complete the training. A correct response pattern was determined from pilot testing and was defined as the response rate within the 3000 ms time‒response window from 10–15 key presses. This response rate was then used in the Instrumental task to calculate the number of reward units to be delivered in correct Go trials. During practice trials, positive or negative feedback was presented on the screen, but no reward was delivered. Feedback for incorrect responses during training was specific to the type of error and could indicate 1) anticipation (responding began before the Go signal appeared), 2) over-responding (two or more responses were given after the Go signal disappeared), 3) an incorrect response pace (too slow or too fast, indicating a response rate under 10 or over 15 key presses, respectively; akin to holding the key down without lifting the finger), or 4) an incorrect response (wrong side key pressed).

During the Instrumental task for the small-reward portion conditions, correct responses in the Go trials were rewarded with either one or two reward units, contingent upon the response rate relative to the participants’ baseline. Specifically, a response rate above the baseline was rewarded with two reward units, whereas a response rate between the baseline (inclusive) and up to 20% below it (inclusive) was rewarded with one unit. Trials with response rates more than 20% below the baseline were considered incorrect. Additionally, if participants committed any kind of error listed above (including errors consisting of responses to the NoGo signal), the trial was considered incorrect, and negative feedback (a red cross) was displayed. Withholding a correct response in NoGo trials received only positive feedback without reward.

In the large-reward conditions, correct responses in the Go trials were rewarded with one, two, or three reward units, contingent upon the response rate relative to the participants’ baseline. Specifically, a response rate surpassing the baseline was rewarded with three reward units, whereas a response rate between the baseline (inclusive) and up to 20% below the baseline was rewarded with two units. Trials with response rates between 20% (inclusive) and 40% below the baseline were rewarded with one unit. Response rates below 40% (inclusive) were considered incorrect. For the NoGo trials, and in contrast to the small-reward conditions, correctly withheld responses were rewarded with one piece of reward. In summary, each participant in the large-reward conditions could win a maximum of 584 [mean large reward = 482.94; S.D.= 54.36] pieces of food reward (corresponding to approximately 470 kcal) for the entire experiment, whereas participants in the small-reward conditions could only win a maximum of 384 [mean small reward = 290.91; S.D.= 28.87] reward units (corresponding to approximately 240 kcal). At the end of the Instrumental task, measures of the number of reward units left by the participants were assumed to have indicated that the participant reached saturation, which occurred only in the large-reward condition [mean food left = 61.81; S.D. = 94.58]. Finally, participants in the immediate reward consumption conditions were allowed to consume the reward units they won at the end of each trial, whereas participants in the delayed reward consumption conditions accumulated the reward units and were only allowed to consume them during the pauses between blocks.

The Instrumental task comprised five blocks, each consisting of 40 trials (30 Go trials and 10 NoGo trials), resulting in a total of 200 trials. All 40 trials in each block were presented to the participants in random order, with the only constraint being that no more than 2 NoGo trials could appear consecutively. The participants could win up to 60 (small reward) or 100 (large reward) reward units per block (300 or 500 units in total for the entire task, respectively). The structure of each trial, in both the training phase and the Instrumental task, was the same. Each trial began with a fixation cross displayed for a jittered duration of 1200–1500 ms. Subsequently, a Go or NoGo signal appeared on either the left or right side of the screen for 3000 ms. After that, the 60 × 60 mm white square remained visible for an additional 700 ms. All stimuli were depicted on a screen subtending approximately 10.4 degrees of the visual angle.

The participants were instructed to to press the C or M keys on the computer keyboard with their left or right index finger, respectively, depending on the side where the Go stimulus was presented on the screen. They were also told to withhold their response in the trials where a NoGo signal was presented. Feedback was provided 1000 ms after the white square disappeared for 300 ms. Following a correct Go trial, the feedback presented consisted of an image of the reward units won. A positive feedback symbol (a green checkmark) was presented in correct NoGo trials (small-reward condition). The reward was always delivered simultaneously with the feedback presentation in each rewarded trial across the task.

##### Transfer task

The final task of the experiment involved a transfer phase that incorporated an extinction procedure. This last task was similar to the previous Instrumental task but differed in two key aspects: i) the Go and NoGo signals were preceded by either the CS+ or a single CS- compound from the initial Pavlovian task, and ii) the ratio of Go to NoGo trials was now equal (50%). The extinction procedure involved providing uninformative feedback for each trial and withholding reward delivery during the whole task. Consequently, the feedback presented at the end of each trial always corresponded to an interrogative sign, regardless of the participants’ response. The task instructions mirrored those of the Instrumental task, except that the participants were informed that any rewards earned would be delivered at the end of the task. This procedure, often referred to as nominal extinction, has been widely adopted to keep participants engaged during the transfer task without further reinforcing the learned association (Allman et al., 2010; Hogarth & Chase, 2011; Prévost et al., 2012; Talmi et al., 2008; Watson et al., 2014). The participants were also instructed to disregard the CS compounds and to respond as they did in the preceding Instrumental task. The structure of stimulus presentation within each trial was identical to those in the previous two tasks combined (see Figure 1C).

The transfer task comprised five blocks composed of 40 trials (200 trials in total), with 20 trials corresponding to the Go condition and 20 to the NoGo condition in each block. The 40 trials in each block were presented to the participants in random order, without any restrictions or constraints. Additionally, only one of the 3 CS- compounds was used in this task, which corresponded always to the most different one in relation to the CS+ compound, i.e., the one composed of a different image and a most different tone. All the stimuli on the screen subtended approximately 10.4 degrees of the visual angle.

#### Statistical analysis

First, direct comparisons were performed to evaluate differences in *blood glucose concentrations, hunger feelings, explicit liking*, and *explicit wanting* before and after the tasks. For the learning test in the Pavlovian task, a linear mixed model with a binary response variable was built to predict whether participants would have explicitly learned the contingencies between CS compounds and reward delivery during the task. The model included only the intercept term, as well as the between-subjects and/or the *CS-type* factors, if they improved the models’ fit. Next, performance in the different phases of the experiment was tested in separate mixed-design ANOVA models, which included two between-subjects factors (*reward portion* and *reward timing*). For the response block of the Pavlovian task, the dependent variables of the models were *response latency* and *pedal presses*, and the within-subjects factor was *CS type* (CS+ vs. CS-). For *pedal presses*, weighted percentatges for each *CS type* and participant were computed to ensure that each trial was properly accounted for, given that there were unequal numbers of trials across conditions. For the Instrumental task, the dependent variable of the model corresponded to the *response rate* (number of key presses in Go trials), and the within-subjects factor was *block* (5 levels: blocks 1–5). Moreover, for the transfer task analysis, all ANOVA models were built by subtracting differences in *response rate* and *response latency* for *CS types* (CS+ minus CS-) and then using these values as the dependent variables to test reward devaluation. Finally, for the analysis of performance on NoGo trials, the dependent variable was the percentage of correct response withholdings. The Greenhouse–Geiser correction was applied in all the ANOVAs to correct for violations of the sphericity assumption when necessary. The effect sizes are reported as Cohen’s *d* values for *t tests* and as partial eta-squared values for ANOVAs.

### Results

#### Preliminary measurements

A summary of the results is provided in Table 1. First, we analyzed participants’ *blood glucose* levels before and after the task. The results revealed a main effect of *reward portion* [large reward *Δ* = −29.21, S.D. = 22.5; small reward *Δ* = −2.53, S.D. = 10.14; *F*(1,95) = 59.53; *p* < 0.001; η*_p_*^2^ = 0.39], with no effect of *reward time* [*F*(1,95) = 2.11; *p* = 0.149; η*_p_*^2^ = 0.02]. The interaction term was also nonsignificant [*F*(1,95) = 0.96; *p* = 0.329; η*_p_*^2^ = 0.01], indicating that differences in *blood glucose pre- and postconcentration* corresponded well with our manipulation of food satiation. Similarly, the *subjective hunger* ratings significantly increased in the pre–post decrease in the *large-reward food portion condition* [large reward *Δ* = −3.81, S.D. = 3.16; small reward *Δ* = −0.42, S.D. = 1.32; *F*(1,95) = 48.72; *p* < 0.001; η*_p_*^2^ = 0.34], *reward time* [*F*(1,95) = 0.002; *p* = 0.961; η*_p_*^2^ < 0.001], and *reward portion* x *reward time* [*F*(1,95) = 0.61; *p* = 0.435; η*_p_*^2^ = 0.006], further confirming that the satiation manipulation was successful.

**Table 1.** Preliminary measures (Experiment 1). Mean (S.D.) pre- and post-experiment glucose concentrations, subjective hunger ratings, and explicit wanting scores, reported for each condition and both experiments.

| Condition | Glucose Levels |  | Subjective Hunger |  | Wanting |  |
| --- | --- | --- | --- | --- | --- | --- |
|  | Pre | Post | Pre | Post | Pre | Post |
| LRI | 94.9 (8.1) | 128.3 (26.5) | 7.2 (1.7) | 3.2 (2.8) | 7.8 (1.7) | 1.9 (2) |
| LRD | 94.9 (9.6) | 119.9 (17.4) | 6.4 (2) | 2.8 (2.8) | 7.2 (1.9) | 2.4 (2.1) |
| SRI | 93.8 (6.6) | 97.1 (10.4) | 7.3 (1.9) | 7 (1.4) | 7.8 (2.5) | 5.4 (3.1) |
| SRD | 93.5 (7.5) | 95.2 (13.1) | 7.1 (2.1) | 6.5 (1.7) | 7.6 (2.4) | 4.7 (2.7) |
| Money | 90.5 (9.2) | 84.5 (5.5) | 6.6 (2.2) | 8.3 (1.7) | 7.8 (1.8) | 7.7 (1.8) |
| Food | 94 (6.7) | 97.6 (1) | 7.3 (1.9) | 7 (1.4) | 7.8 (2.6) | 5.5 (3.1) |

Next, the results showed that reward *liking* was rated similarly across all conditions [*reward portion*: *F*(1,95) = 1.65; *p* = 0.202; η*_p_*^2^ = 0.02; *reward time*: [*F*(1,95) = 2.22; *p* = 0.139; η*_p_*^2^ = 0.02; *reward portion* × *reward time*: *F*(1,95) = 0.47; *p* = 0.495; η*_p_*^2^ = 0.004]. For food *wanting*, the results revealed a main effect of *reward portion* [large reward *Δ* = −5.37, S.D. = 2.23; small reward *Δ* = −2.66, S.D. = 3.38; *F*(1,95) = 21.93; *p* < 0.001; η*_p_*^2^ = 0.19], whereas *reward time* [*F*(1,95) = 0.39; *p* = 0.535; η*_p_*^2^ = 0.004] and the interaction term *reward portion* × *reward time* [*F*(1,95) = 1.66; *p* = 0.200; η*_p_*^2^ = 0.02] did not reach statistical significance, indicating that the *explicit wanting* decreased more when participants received large-reward food portions than when they received small-reward portions.

#### Pavlovian learning task

In the explicit learning test, participants responded correctly to the discrimination between CS- and CS+ in 81.31% of the trials. Furthermore, all the participants correctly identified the rewarded image from the CS+ compound and identified the complete audio-visual CS+ in at least 1 of the 2 trials. The participants performed similarly across all the conditions (all *p values >* 0.100). Results from the generalized linear model revealed that the intercept was highly significant (*β* = 1.92, *SE* = 0.106, *z* = 18.02, *p* < 0.001; odds ratio index = 6.76). Since an odds ratio of 1 would be expected by chance (see, for instance, McHugh, 2009), these results suggest good discrimination between CS+ and CS- stimuli.

We subsequently analyzed the performance during the last block of the Pavlovian task (see Table 2 and Figure S1), where participants could either voluntarily press a pedal to reveal the feedback or wait 3500 ms. Response latencies and pedal presses (percentatge of CS+ / CS-) served us to obtain indirect measures of CS+ learning. The results revealed a significant main effect of *CS type* [*F*(1,34) = 10.82; *p* < 0.01; η*_p_*^2^ = 0.24] on response latency, with participants responding faster [CS+ mean = 685.34, S.D. = 224.21; CS- mean = 755.10, S.D. = 159.12] to uncover the reward associated with the CS+ compound. In contrast, no significant effects were observed for *reward time* [*F*(1,34) = 2.03; *p* = 0.163; η*_p_*^2^ = 0.06] or *reward portion* [*F*(1,34) = 1.47; *p* = 0.234; η*_p_*^2^ = 0.04]. Moreover, none of the interaction terms reached statistical significance (all *p values >* 0.150).

**Table 2.**
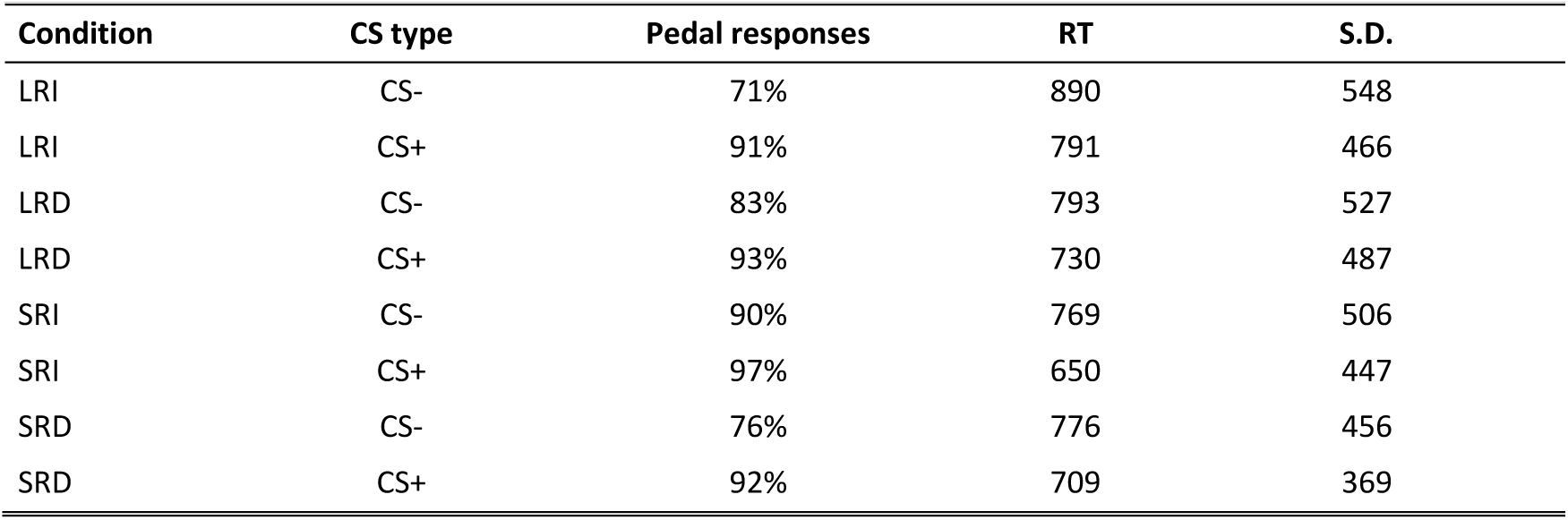
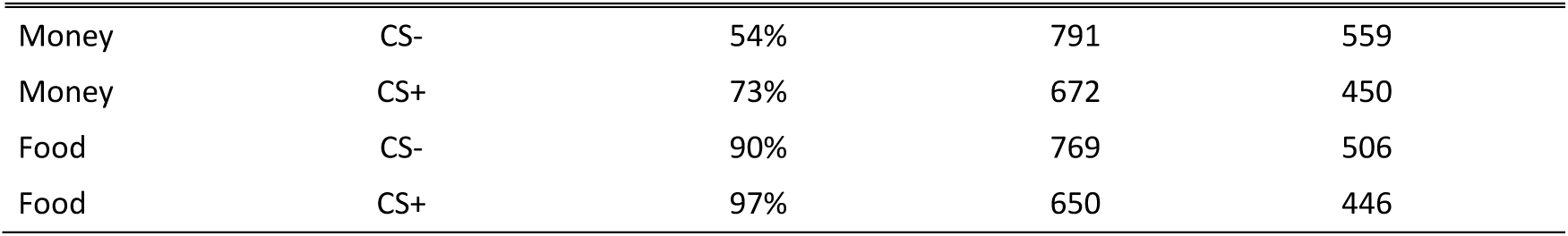
Summary of pedal presses and latencies during the last block of the Pavlovian task). Mean percentage of voluntary pedal responses, response times (RT), and standard deviations (S.D.) for CS+ and CS− compounds across all conditions and both experiments.

| Condition | CS type | Pedal responses | RT | S.D. |
| --- | --- | --- | --- | --- |
| LRI | CS- | 71% | 890 | 548 |
| LRI | CS+ | 91% | 791 | 466 |
| LRD | CS- | 83% | 793 | 527 |
| LRD | CS+ | 93% | 730 | 487 |
| SRI | CS- | 90% | 769 | 506 |
| SRI | CS+ | 97% | 650 | 447 |
| SRD | CS- | 76% | 776 | 456 |
| SRD | CS+ | 92% | 709 | 369 |
| Money | CS- | 54% | 791 | 559 |
| Money | CS+ | 73% | 672 | 450 |
| Food | CS- | 90% | 769 | 506 |
| Food | CS+ | 97% | 650 | 446 |

In terms of voluntary Pedal responses (percentatge of pedal presses), a significant main effect of *CS type* was observed [*F*(1,95) = 26.44; *p* < 0.001; η*_p_*^2^ = 0.22], indicating that participants uncovered the CS+ compound significantly more frequently (93.5%) than the CS− compound (80.2%). No main effects were found for *reward portion* [*F*(1,95) = 1.15; *p* = 0.287; η*_p_*^2^ = 0.01] or *reward time* [*F*(1,95) = 0.05; *p* = 0.818; η*_p_*^2^ = 0.0004]. However, the interaction *reward portion* × *reward time* [*F*(1,95) = 3.98; *p* = 0.048; η*_p_*^2^ = 0.04] reached statistical significance. Further Tukey-adjusted pairwise comparisons revealed significant differences in voluntary responses between the large- and small-reward conditions when the food reward was immediately delivered [large reward = 81%, S.D. = 4.3; small reward = 94%, S.D. = 2.4; *t*(95) = 2.16, *p* = 0.03, *d* = 0.52]. Thus, participants in the SRI condition exhibited a greater percentage of responses to uncover reward information in every trial. The remaining interaction terms were not significant (all *p values* > 0.100).

#### Instrumental learning task

The participants achieved an overall accuracy (correct Go trials and correct inhibitions in NoGo trials) of 88.56% (S.D. = 3.2%) on this task (see Table 3 and Figure S2 for a summary). We observed a significant main effect of *block* for response rates (number of key presses) in correctly rewarded trials [*F*(4,380) = 23.45; *p* < 0.001; η*_p_*^2^ = 0.20], indicating that response rates increased across blocks. A significant main effect of *reward portion* was also observed [mean large reward = 13.01, S.D. = 2.2; small reward = 13.89, S.D. = 1.9; *F*(1,95) = 8.16; *p* < 0.01; η*_p_*^2^ = 0.08], with participants in the small-reward condition responding with a greater number of key presses than those in the large-reward condition. No significant effects of *reward time* were found [*F*(1,95) = 1.33; *p* = 0.251; η*_p_*^2^ = 0.01], and all the interaction terms were nonsignificant (all *p values* > 0.3; see Figure 2).

**Figure 2.**
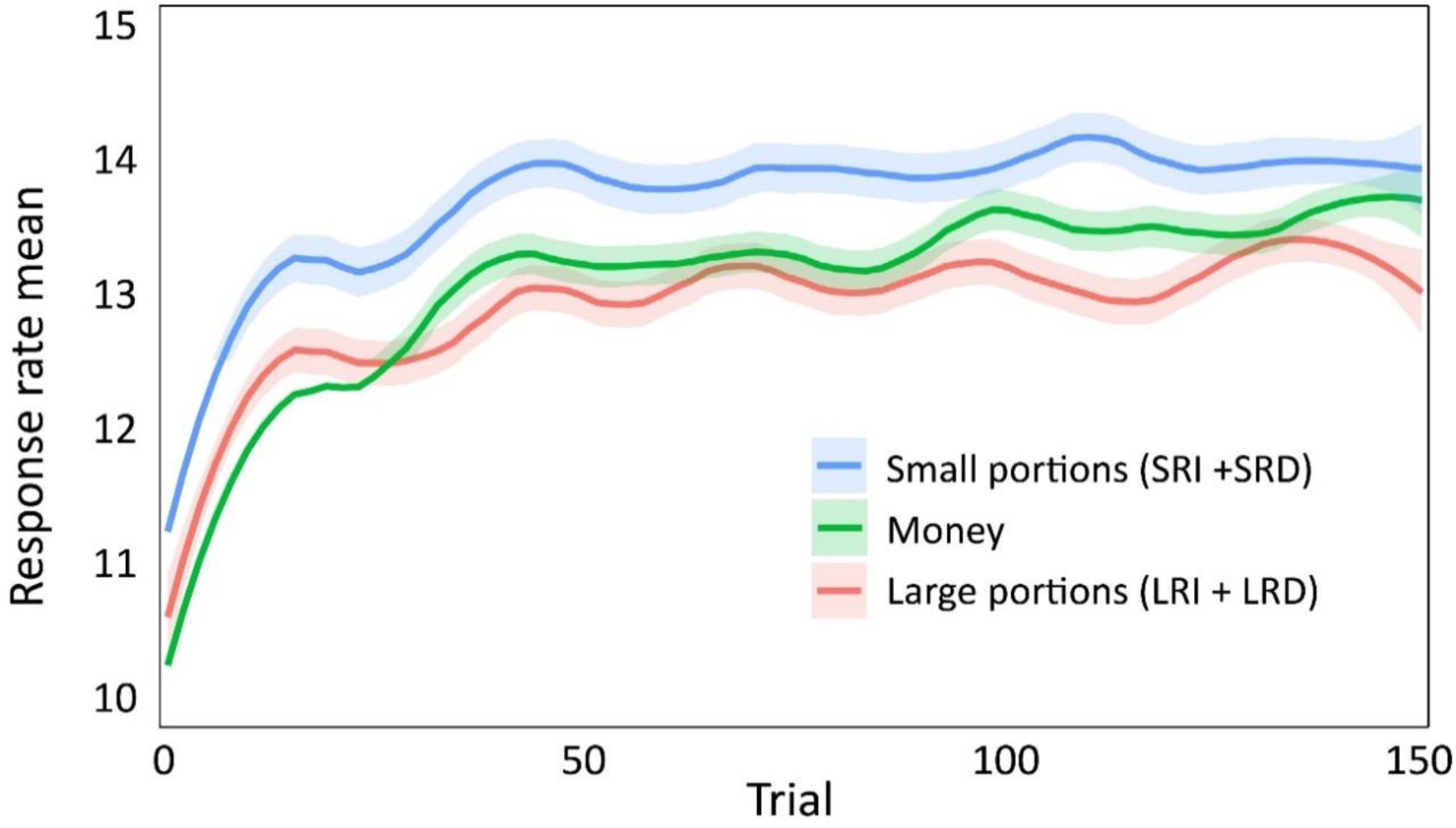
Mean numbers of keys pressed (response rates) among trials during the Instrumental task in both experiments. The *response rate* (number of key presses) was averaged by the *reward portion* (large and small). The immediate and delay conditions were collapsed because they did not yield any effect. Note that trials with any error were considered incorrect and were not included.

**Table 3.** Instrumental task performance. Mean number of key presses (S.D.) during the Instrumental task, reported by block and condition, from Experiment 1 and Experiment 2.

| Block | SRD | SRI | LRD | LRI | Money | Food |
| --- | --- | --- | --- | --- | --- | --- |
| 1 | 12.9 (1.2) | 13.5 (1.8) | 12.2 (1.8) | 12.6 (1.4) | 12.2 (1.8) | 13.5 (1.8) |
| 2 | 13.6 (1.4) | 14.3 (1.7) | 13 (2) | 13 (1.7) | 13.1 (2.2) | 14.3 (1.7) |
| 3 | 13.7 (1.4) | 14.4 (1.4) | 13.1 (1.9) | 13.3 (1.9) | 13.2 (2) | 14.4 (1.4) |
| 4 | 13.9 (1.4) | 14.4 (1.5) | 13.3 (2) | 13.3 (1.7) | 13.5 (2) | 14.4 (1.5) |
| 5 | 13.8 (1.3) | 14.3 (1.6) | 13.5 (1.7) | 13.3 (1.7) | 13.6 (1.8) | 14.3 (1.6) |

Finally, the results of the participants’ performance in the NoGo trials revealed no significant main effect of *reward portion* [*F*(1,95) = 1.83; *p* = 0.179; η*_p_*^2^ < 0.02]*, block* [*F*(4,380) = 1.76; *p* = 0.143; η*_p_*^2^ = 0.02], or *reward time* [*F*(1,95) = 0.33; *p* = 0.564; η*_p_*^2^ = 0.02]. Moreover, none of the interaction terms reached statistical significance (all *p values* > 0.400). Thus, participants’ performance in NoGo trials remained consistent across all experimental conditions.

#### Transfer task

Next, participants’ performance in the transfer task of the experiment was analyzed (see Figure 3A and S3, and Tables 4 and 5). The participants’ overall performance (correct Go trials and correct inhibitions in NoGo trials) in this task was 87.3% (S.D. = 3.3%), confirming its successful implementation. The ANOVA on response rates (number of key presses), when considering all blocks, revealed nonsignificant main effects of *reward time* [*F*(1,95) = 0.57; *p* = 0.453; η*_p_*^2^ = 0.006], *reward portion* [*F*(1,95) = 0.15; *p* = 0.903; η*_p_*^2^ = 0.0004], or *block* [*F*(4,380) = 1.61; *p* = 0.178; η*_p_*^2^ = 0.02]. Moreover, none of the interaction terms were statistically significant (all *p* values > 0.19). However, when the analysis was limited to the first two blocks—when the transfer effect was most pronounced (see, e.g., Talmi et al., 2008)—the results for response rates revealed a significant main effect of *reward time* [immediate reward *Δ* = 0.44, S.D. = 1.12; delayed reward *Δ* = 0.21, S.D. = 1.01; *F*(1,95) = 4.36; *p* = 0.039; η*_p_*^2^ = 0.04], with no significant difference observed for *block* [*F*(1,95) = 0.01; *p* = 0.920; η*_p_*^2^ = 0.0001] or *reward portion* [*F*(1,95) = 1.21; *p* = 0.274; η*_p_*^2^ = 0.01]. However, the interaction term *reward portion* × *block* reached statistical significance [*F*(1,95) = 4.4; *p* = 0.039; η*_p_*^2^ = 0.05]. Tukey-adjusted pairwise comparisons revealed a marginally significant difference between the *reward portion* conditions in the second block [mean large reward *Δ* = 0.08, S.D. = 0.6; small reward *Δ* = 0.48, S.D. = 1.32; *t*(95) = 1.92; *p* = 0.058; *d* = 0.38], suggesting greater food reward devaluation for participants in the large-reward condition (Figure 3A). Finally, the three-way interaction *reward time* × *reward portion* × *block* showed a trend toward significance [*F*(1,95) = 3.70; *p* = 0.057; η*_p_*^2^ = 0.04], whereas the remaining interaction terms were not statistically significant (all *p values* > 0.4).

**Figure 3.**
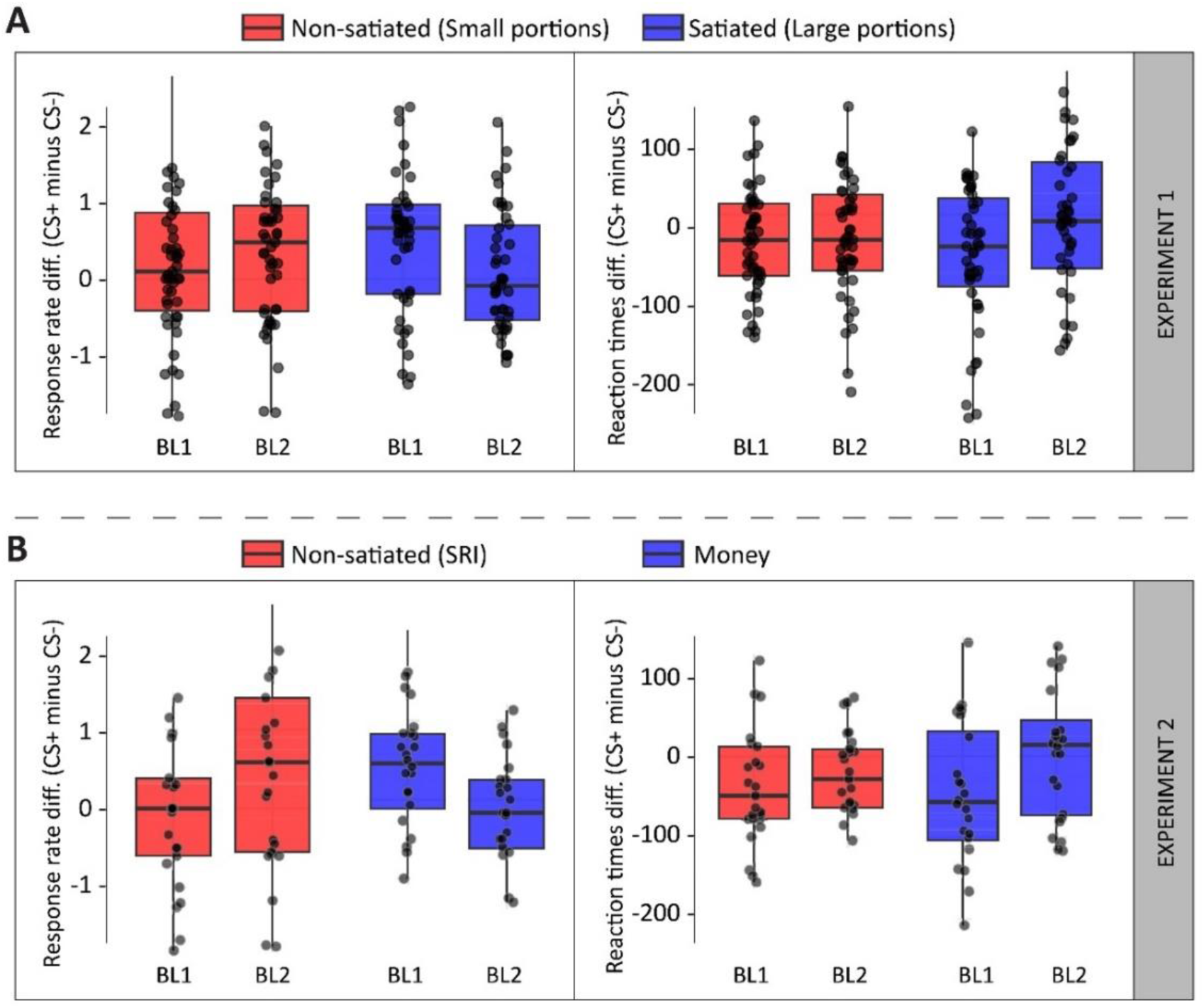
Interaction effects of *reward portion* × *block* on *response rate* and *response latency* (in milliseconds) differences (CS+ minus CS−) from the two experiments. The figure illustrates the interaction between the *reward portion* and *block* conditions during the transfer task of Experiment 1 (**A**), as well as the interaction between the *food* and *money* conditions during the same task of Experiment 2 (**B**).

**Table 4.** Response rate during the Transfer task. Mean number of key presses (S.D.) across all conditions, both experiments, and the two CS types.

| Block | CS type | LRI | LRD | SRI | SRD | Money | Food |
| --- | --- | --- | --- | --- | --- | --- | --- |
| 1 | CS- | 12.45 (2.4) | 13.24 (2.3) | 13.53 (2.0) | 13.20 (1.7) | 12.96 (2.1) | 13.53 (2.0) |
| 1 | CS+ | 13.04 (2.3) | 13.32 (2.1) | 13.69 (2.1) | 13.34 (1.7) | 13.24 (1.9) | 13.69 (2.1) |
| 2 | CS- | 12.45 (2.4) | 13.11 (2.2) | 13.37 (2.4) | 13.11 (1.7) | 13.16 (1.9) | 13.49 (2.1) |
| 2 | CS+ | 12.69 (2.4) | 13.11 (2.1) | 13.83 (2.3) | 13.25 (1.8) | 13.12 (2) | 13.83 (2.3) |

**Table 5.** Response latencies during the Transfer task. Mean reaction times (latency from the first key press; S.D. in parentheses) across all conditions, both experiments, and the two CS types.

| Block | CS type | LRI | LRD | SRI | SRD | Money | Food |
| --- | --- | --- | --- | --- | --- | --- | --- |
| 1 | CS- | 622 (170) | 527 (77) | 539 (79) | 549 (129) | 534 (190) | 539 (79) |
| 1 | CS+ | 549 (164) | 506 (96) | 495 (72) | 512 (129) | 478 (114) | 495 (72) |
| 2 | CS- | 600 (174) | 533 (104) | 556 (108) | 546 (128) | 499 (143) | 556 (108) |
| 2 | CS+ | 596 (186) | 541 (130) | 494 (84) | 524 (125) | 493 (132) | 494 (84) |

The response latencies during the extinction procedure were subsequently analyzed (Table 5). When all the blocks were considered together, no significant main effects of *reward time* [*F*(1,95) = 3.31; *p* = 0.072; η*_p_*^2^ = 0.03], *reward portion* [*F*(1,95) = 2.19; *p* = 0.142; η*_p_*^2^ = 0.02], or *block* were observed [*F*(4,380) = 2.08; *p* = 0.099; η*_p_*^2^ = 0.02]. Moreover, none of the interaction terms were statistically significant (all *p* values > 0.200). However, when only the first two blocks were entered into the analysis, the results revealed a trend toward significance for a main effect of *reward time* [*F*(1,95) = 3.5; *p* = 0.064; η*_p_*^2^ = 0.04]. In contrast, no significant main effect of *reward portion* was observed [*F*(1,95) = 1.5; *p* = 0.224; η*_p_*^2^ = 0.02], although the interaction term *reward portion* × *block* reached statistical significance [*F*(1,95) = 4.49; *p* = 0.037; η_p_ ^2^ = 0.05]. The main effect of *block* also reached statistical significance [*F*(1,95) = 3.98; *p* = 0.049; η*_p_*^2^ = 0.04]. Post hoc comparisons revealed that differences in response latencies between *reward portions* were more pronounced in the second block [mean large reward *Δ* = 1.41, S.D. = 97.1; small reward *Δ* = −40.7, S.D. = 95.8; *t*(95) = −2.18; *p* = 0.031; *d* = 0.44] and that participants in the small-reward condition were more resistant to devaluing the CS+ contingencies (see Figure 3A). The remaining interaction terms were not statistically significant (all *p values* > 0.1).

Participants’ performance in NoGo trials was also analyzed. The results revealed significant main effects of *block* [*F*(4,380) = 3.04; *p* = 0.021; η*_p_*^2^ = 0.03] and *CS type* [*F*(1,95) = 11.07; *p* < 0.01; η*_p_*^2^ = 0.11], as well as a significant interaction between *block* × *CS type* [*F*(4,380) = 3.39; *p* = 0.012; η*_p_*^2^ = 0.03]. Further post hoc comparisons revealed significant differences between *CS types* only in the first block [mean CS+ = 94.3%, S.D. = 8.4; CS- = 97.7%, S.D. = 4.4; *t*(95) = 3.75; *p* < 0.001; *d* = 0.5], indicating that participants committed more errors in NoGo trials during the first block, particularly in CS- trials. A nonsignificant main effect of *reward portion* [*F*(1,95) = 0.20; *p* = 0.655; η*_p_*^2^ = 0.002] or *reward time* [*F*(1,95) = 0.37; *p* = 0.545; η*_p_*^2^ = 0.003] was observed. The remaining interaction terms were not statistically significant (all *p values* > 0.1).

### Discussion

During the Pavlovian phase, participants acquired the cue‒food association, as indicated by faster response latencies and a higher frequency of pedal presses for the reward-associated compound in the last block using pedal responses to discover the feedback. Although reward consumption timing did not significantly influence these learning indices, the SRI condition yielded a higher percentage of pedal presses. This pattern implies that frequent, smaller reward consumption can sustain engagement in cue-driven tasks, as revealed by the interaction found between reward portion and reward timing. This result is in line with the findings from animal studies showing that responsiveness to food cues persists until satiety is reached (Panayi & Killcross, 2022). Participants receiving small rewards maintained elevated response rates throughout the Instrumental phase. This outcome is consistent with the notion that smaller portions delay the onset of satiety, thereby prolonging reward-seeking behavior (Lingawi et al., 2022).

The absence of a main effect of reward consumption timing on increased instrumental response rates further suggests that immediate versus delayed consumption does not substantially influence action rates when portion size is the critical factor modulating satiety (Marshall & Ostlund, 2021). During the transfer phase, the early blocks revealed that participants who received larger, immediate rewards exhibited faster declines in cue-elicited responding—a behavioral pattern consistent with rapid devaluation driven by satiation (Burghoorn et al., 2024). Conversely, participants in the small-reward conditions appeared more resistant to devaluation, likely because the modest portions did not elicit satiety-related responses. These findings align with our initial hypothesis that large, immediately consumed rewards result in a faster reduction in instrumental responding, whereas small or delayed reward consumption helps maintain behavioral engagement.

Finally, the results concerning hunger feelings and blood glucose levels indicated that manipulating portion size effectively induced within-session satiety under large-reward conditions (Ello-Martin et al., 2005). In contrast, immediate versus delayed consumption, when considered in isolation, did not significantly alter these physiological or subjective measures, suggesting that portion size was the primary driver of satiety in the current experiment (Ueland et al., 2009).

In summary, our data support the idea that real-time ingestion of palatable foods results in measurable, within-session shifts in motivation and reward-seeking behavior (Aitken et al., 2016). Although delayed consumption alone may not yield pronounced satiety-related declines, its combination with larger portions appears to modulate the rate at which participants reduce their food-seeking behavior (Sommer et al., 2022). These outcomes underscore the importance of capturing natural hunger–satiety cycles in human PIT studies to better replicate the conditions observed in animal models (Colagiuri & Lovibond, 2015).

Building on these results, we designed a second experiment to further examine whether the encountered devaluation effects are specific to food (primary rewards) or extend to other reinforcers, such as monetary gains (secondary rewards). Thus, in Experiment 2, immediate edible rewards were directly compared with real-time monetary payouts while employing only small food portions to minimize satiety effects. This approach allowed us to assess whether the mechanisms underlying cue-driven behavior generalize to other reward types that do not elicit physiological satiety, such as monetary gains.

## Experiment 2

Although money is often described as a universal reinforcer, few studies have directly compared food and monetary rewards under conditions that mirror real-time delivery in animal models (Lehner et al., 2017). In Experiment 2, we compare two reward types delivered incrementally in a similar manner: small food portions that are consumed on a trial-by-trial basis versus monetary rewards that are earned in the same manner but not immediately used in any expenditure (Garofalo & di Pellegrino, 2017). Moreover, by using only small food portions—thus avoiding the satiety effects observed with larger food portions in Experiment 1—we aim to directly compare the two reward types under equivalent delivery and satiety conditions (Colagiuri & Lovibond, 2015).

A new group of participants underwent the three PIT phases again, with rewards delivered in real time (money in this case). In the food condition (SRI condition from Experiment 1), participants consumed a small portion of food in each trial, whereas in the monetary condition, they earned money (small amount of units and immediate delivery, matching the food condition) with no concomitant consumption. This design allowed us to assess whether cue-driven behavior is modulated by real-time reinforcement when the reward neither induces satiety nor provides immediate gratification (Hogarth & Chase, 2011).

We hypothesize that if the mechanisms underlying cue-driven behavior operate similarly regardless of reward type, both conditions will show parallel patterns of learning, transfer, and devaluation. Alternatively, if monetary rewards are inherently less prone to devaluation—due to the absence of any hunger–satiety modulation—we may observe more persistent instrumental responding and less decline in cue-elicited behavior than in the food condition. As such, Experiment 2 served as a control experiment to compare the dynamics of food versus monetary rewards, thereby shedding light on the interplay between satiety-independent valuation and the processes by which cues from the two reward types invigorate behavioral output.

### Methods

#### Participants

Data from participants in the small and immediate reward portion conditions of Experiment 1 are included here solely for comparison. Hence, 29 students (23 female) from the Faculty of Psychology at the University of Barcelona, who did not take part in Experiment 1, participated in this experiment [mean age = 19.48 years, S.D. = 1.7, range = 17–24]. A power analysis using G*Power (Erdfelder et al., 2009) indicated that a sample of 46 participants (23 in each group) was sufficient to assess the effect of 2 within-subjects variables and 1 between-subjects variable (with a power = 95% and an a priori alpha set at p = 0.05) for a moderate effect size estimate (η*_p_*^2^ = 0.06). All participants had BMIs within the normal range (18–25) [mean BMI, food-reward condition = 21.76, S.D. = 2.3; money-reward condition = 21.87, S.D. = 3.13; *t*(51) = 0.13; *p* = 0.898; *d* = 0.03]. The fasting times (in hours) were similar across conditions [food: 16.66, S.D. = 0.98; money: 16.52, S.D. = 1.12; *t*(51) = 0.51; *p* = 0.619; *d* = 0.14]. All participants provided informed consent as approved by the local ethics committee (IRB00003099) before their participation and received 20€ for completing the experiment.

### Materials & Procedure

#### Preliminary experimental measurements

The procedure for Experiment 2 was identical to that for Experiment 1 (using only small amount and immediate delivery of rewards), except for the use of a monetary reward (5¢ coins), in the money condition, instead of food. As in Experiment 1, the participants were asked first to evaluate their subjective measures of *hunger*, *liking* and *wanting* of the reward (food and/or money) used in the experiment. Similarly, when money was used as a reward, participants were first informed about the amount of 5¢ coins and the total amount of money they could earn during the experiment. They then responded to the question *“How much do you like to receive money as a reward during the experiment?”* using the same VAS as in Experiment 1. Finally, the participants were required to report their immediate subjective desire (wanting/craving) to receive (5¢ coins) again using a 10 cm VAS (0 indicated *no desire at all,* and 10 indicated *extreme desire*).

### Results

#### Preliminary measurements

We first analyzed the participants’ pre- task and posttask *blood glucose concentrations* (Table 1). The results revealed a significant decrease in glucose concentration for the *money-reward* condition [money *Δ* = −6.03, S.D. = 5.98; food *Δ* = 3.58, S.D. = 8.96; *t*(51) = −4.66; *p* < 0.001; *d* = −1.29]. This decrease corresponded to the metabolic effect of not consuming approximately 240 kcal. throughout the experiment, as occurred for participants in the food-reward condition. Similarly, the posttask *subjective hunger* level reported by the participants significantly increased compared with the pretask level when money was used as a reward [money *Δ* = 1.75, S.D. = 1.76; food *Δ* = −0.27, S.D. = 1.38; *t*(51) = 4.57; *p* < 0.001; *d* = 1.26]. This finding indicated that the subjective hunger levels in the food reward condition did not change as much as those in the money condition did, indicating that the usual hunger increase that could occur after approximately 3.5 hours (the duration of the entire experiment), considering that the participants accumulated approximately 16.55 hours of fasting upon their arrival at the laboratory.

Next, *explicit liking* and *wanting* of food and money as rewards were analyzed. The results showed that both reward types were rated similarly [mean food = 8.13, S.D. = 1.04; money = 7.48, S.D. = 1.79; *t*(51) = −1.54; *p* = 0.13; *d* = 0.43]. However, when *explicit wanting* was considered, the results revealed that only wanting food as a *reward* decreased throughout the experiment [food *Δ* = −2.35, S.D. = 3.61; money *Δ* = −0.08, S.D. = 1.42; *t*(51) = 3.11; *p* < 0.01; *d* = 0.86].

#### Pavlovian learning task

We began by verifying whether participants learned the audio-visual CS compounds during the Pavlovian task of the PIT experiment. In the explicit learning test, participants correctly differentiated between CS+ and CS- compounds in 83.96% of the trials in the test. Additionally, all participants successfully identified the correct image of the CS+ compound and correctly identified the audio-visual CS+ compound in at least 1 of the 2 trials in which the compound was presented. Furthermore, the results from the generalized linear model revealed that the intercept was highly significant (*β* = 2.290, *SE* = 0.168, *z* = 13.63, *p* < 0.001; odds ratio index = 9.87), indicating that the odds of giving a correct response were approximately 10 times greater than the odds of giving an incorrect response. These results suggest good discrimination between CS+ and CS- stimuli.

We further confirmed the learning of the CS+ compound by analyzing the performance on the last block (Pedal responses) of the Pavlovian task (see Table 2 and Figure S1). The results revealed statistically significant differences in response latencies between *CS types* [*F*(1,14) = 7.12; *p* = 0.018; η*_p_*^2^ = 0.34], indicating a faster response time for the CS+ compound. The main effects of *reward type* [*F*(1,14) = 0.02; *p* = 0.894; η*_p_*^2^ = 0.001] and the interaction term [*F*(1,14) = 1.52; *p* = 0.238; η*_p_*^2^ = 0.10] did not reach statistical significance, suggesting that participants responded in similar ways in both reward-type conditions to uncover the CS+ feedback. In terms of voluntary response percentatge (pedal presses in Both CS types), main effects of *reward type* [*F*(1,51) = 26.18; *p* < 0.001; η*_p_*^2^ = 0.34] and *CS type* were found [*F*(1,51) = 8.37; *p* < 0.01; η*_p_*^2^ = 0.14], denoting that participants in the food *reward* condition responded more often than when receiving money as a reward and that in both *reward types,* all participants responded proportionally more often when presented with a CS+ (84.9%) than when presented with a CS− compound (71.5%). The interaction term did not reach statistical significance [*F*(1,51) = 1.83; *p* = 0.181; η*_p_*^2^ = 0.03].

#### Instrumental learning task

We next analyzed whether participants correctly learned the contingencies between operant responses (key presses) and reward delivery during the Instrumental task (see Table 3 and Figure S2 for a summary). The participants achieved an overall mean response (correct Go trials and correct inhibitions in NoGo trials) accuracy of 88.75% (S.D. = 3.2%). Analysis of the *response rate* in all rewarded trials revealed significant main effects of *block* [*F*(4,204) = 22.52; *p* < 0.001; η*_p_*^2^ = 0.31] and *reward type* [*F*(1,51) = 5.51; *p* = 0.023; η*_p_*^2^ = 0.10], indicating that participants’ response rate increased over time during the task and that food elicited more vigorous response patterns than money when used as a reward. The interaction term was nonsignificant [*F*(4,204) = 1.55; *p* = 0.189; η*_p_*^2^ = 0.03; see Figure 2).

Participants’ performance in NoGo trials was also analyzed. The results of the ANOVA revealed nonsignificant main effects of *reward type* [*F*(1,51) = 0.05; *p* = 0.828; η*_p_*^2^ < 0.01] and *block* [*F*(4,204) = 0.65; *p* = 0.625; η*_p_*^2^ = 0.01]. The interaction term also did not reach statistical significance [*F*(4,204) = 1.91; *p* = 0.110; η*_p_*^2^ = 0.04], indicating that participants’ performance when withholding responses during the task was similar regardless of the *reward type* used.

#### Transfer task

Participants’ performance in the transfer task of the experiment was analyzed to explore the combined effects of the previous Pavlovian and Instrumental tasks and, crucially, to compare the devaluation between the two different reward types employed in this experiment. The participants achieved an accuracy (correct Go trials and correct inhibitions in NoGo trials) of 87.8% (S.D. = 3.3%) in the task (see Figure 3B and S3, and Tables 4 and 5). The results for *response rates* revealed no main effect of *reward type* [*F*(1,50) = 2.27; *p* = 0.139; η*_p_*^2^ = 0.05] or *block* [*F*(1,50) = 0.64; *p* = 0.429; η*_p_*^2^ = 0.01], although the interaction term *reward type* × *block* was found to be statistically significant [*F*(1,50) = 5.70; *p* = 0.021; η*_p_*^2^ = 0.10]. Post hoc comparisons revealed that differences in *response rates* between *CS types* were similar for both *reward types* in *block* one [*t*(50) = 0.23, *p* = 0.821, *d* = 0.06]. However, in *block* two, a significant reduction was observed for participants in the money-reward condition [mean money *Δ* = 0.03, S.D. = 0.4; food *Δ* = 0.80, S.D. = 1.2; *t*(50) = −2.19, *p* = 0.03, *d* = 0.61], suggesting that food was more resistant to devaluation than money when participants were not satiated.

A subsequent analysis of response latencies yielded comparable results (Table 5). There were no main effects of *reward type* [*F*(1,50) = 2.51; *p* = 0.149; η*_p_*^2^ = 0.04] or *block* [*F*(1,50) = 1.14; *p* = 0.291; η*_p_*^2^ = 0.02], although the interaction term *reward type* × *block* reached statistical significance [*F*(1,50) = 5.85; *p* = 0.019; η*_p_*^2^ = 0.10]. Subsequent Tukey-adjusted direct comparisons revealed that response latencies for both *reward types* were similar in *block* one [*t*(50) = −0.48, *p* = 0.628, *d* = −0.12]. In *block* two, however, a significant reduction in Δ response latency was observed in the money-reward condition [mean Money *Δ* CS = −6.63, S.D. = 72.7; mean Food *Δ* CS = −53.78, S.D. = 82.2; *t*(50) = 2.43, *p* = 0.019, *d* = 0.61], which is consistent with the response rate results, suggesting that when food was used as a reward, it was more resistant to devaluation than money was in the current study (Figure 3B).

Finally, participants’ performance on NoGo trials was also analyzed. The results revealed significant main effects of *block* [*F*(4,204) = 2.66; *p* = 0.033; η*_p_*^2^ = 0.05] and *CS type* [*F*(1,51) = 4.53; *p* = 0.038; η*_p_*^2^ = 0.08], indicating that the error rate decreased as the task progressed and that errors were more frequent when participants were presented with the CS+. The main effect of *reward type* [*F*(1,51) = 0.06; *p* = 0.802; η*_p_*^2^ = 0.001] did not reach statistical significance. Moreover, none of the interaction terms were statistically significant (all *p* values > 0.2).

### Discussion

Experiment 2 was designed as a control experiment to compare the effects of two reward types—small, immediately consumed food rewards versus real-time monetary rewards—on Pavlovian and Instrumental learning and their subsequent devaluation during an extinction procedure. Our findings indicate that both reward types facilitate initial cue-driven learning, but subtle differences emerge in the persistence of response. Hence, participants in the food condition presented slightly higher response rates during both the Pavlovian and Instrumental phases and maintained their responses more consistently during the early blocks of the transfer phase. In contrast, those receiving monetary rewards experienced a more rapid decline in performance by the second transfer block.

These results suggest that although monetary rewards are effective in generating cue-driven behavior, their motivational value may decrease more quickly than food rewards do (Lehner et al., 2017). This difference likely reflects the absence of physiological signals in monetary rewards (Garofalo & di Pellegrino, 2017). Although food rewards engage in biological feedback mechanisms associated with hunger and satiety—which may help sustain their motivational value over time—the nonconsumable nature of money indicates that its value is primarily subject to cognitive or contextual factors (Colagiuri & Lovibond, 2015).

Thus, although both reward types initially invigorate behavior, food appears to retain its influence for slightly longer when delivered in small, real-time portions for immediate consumption, thereby avoiding satiation. These findings contribute to our understanding of how hunger/satiety-independent valuation interacts with cue-driven processes and provide a basis for comparing reward types in real-time reward delivery studies.

## General discussion

Overall, our findings indicate that in humans, Pavlovian cues can strongly affect motivation for eating behavior, and this effect persists even when the outcome loses its initial value due to satiety measured under extinction. The results of Experiment 1 revealed that although satiety accelerates the decline in cue-driven behavior, it does not fully eliminate it. Thus, participants receiving large, immediately consumable portions reached satiety quickly, which led them to reduce instrumental responses at a faster rate than those receiving smaller, more frequent portions. Moreover, the results of Experiment 2 revealed that although cues for monetary rewards also increase eating behavior, these secondary incentives lose their motivational impact more quickly than does food under nonsatiated conditions. Taken together, these findings highlight a central observation: satiety is not required for cue-induced effects on eating to diminish, but it modulates the timing of the behavioral shift by accelerating outcome devaluation (Rescorla, 1994; Watson et al., 2014).

These findings align with ideas from classic rodent paradigms, where repeated within-session consumption of tangible rewards naturally leads to reward devaluation via satiety (Balleine & Dickinson, 1998; Epstein et al., 2010; Rolls et al., 2007). In the present study, human participants receiving large, immediately consumed portions showed accelerated devaluation—a sharper and faster decline in instrumental responding once satiety was reached—than did those receiving smaller portions that did not induce satiety. This outcome contrasts with earlier human PIT experiments that predominantly used symbolic reinforcers or hypothetical outcomes and therefore lacked physiological feedback (Eder & Dignath, 2016; Hogarth & Chase, 2011; Kral & Rolls, 2004; Rolls et al., 2007; Watson et al., 2014; Zlatevska et al., 2014).

Our PIT approach, which involves delivering tangible food on a trial-by-trial basis and systematically manipulating both portion size and consumption timing, more closely mirrors rodent paradigms (Corbit & Balleine, 2005; Holland, 2004). In these paradigms, repeated small- pellet deliveries allow for the gradual tracking of hunger–satiety fluctuations, with robust declines in lever-pressing once animals reach satiety within sessions (Epstein et al., 2010; Panayi & Killcross, 2022). Interestingly, during the Pavlovian and Instrumental phases of our study, portion size did not directly affect participants’ ability to learn cue–reward contingencies or response–outcome mappings until the end of the Instrumental task, when satiation emerged in the large portions’ conditions. These data align with findings from Colagiuri & Lovibond (2015), who showed that early-stage associative learning remains robust across conditions. However, during the transfer phase, large, immediately consumed portions resulted in faster behavioral decline as participants experienced real-time satiety, echoing the devaluation thresholds described in animal studies (Epstein et al., 2010; Rolls et al., 2007). Conversely, smaller portions prevented satiety build-up, leading to prolonged responding and aligning with previous studies in which partitioned intakes were found to induce approaching behavior and delay devaluation (Zlatevska et al., 2014).

The main difference between our design and other human PIT tasks lies in the immediacy and actuality of food intake—precisely, the condition that rodent studies use to detect clear within-session satiation (Holland, 2004; Panayi & Killcross, 2022). Interestingly, once satiety was reached, participants adjusted their effort, demonstrating flexibility in response devaluation (Lingawi et al., 2022; Tricomi et al., 2009). Ultimately, our findings suggest that by integrating immediate consumption, portion size variation, and real food rewards, human PIT studies are able to replicate the dynamic devaluation patterns observed in rodent models and better capture the natural regulation of cue-driven behaviors (Holland, 2004; Pool et al., 2016).

Another interesting finding emerged in the delayed consumption conditions in Experiment 1, where participants accumulated rewards until the end of each block before consuming them. In animal studies, postponing an outcome can prolong goal-directed performance under certain conditions (Urcelay & Jonkman, 2019), either because satiation remains unreached or because the reward retains its appeal. In the present study, participants who had to wait before ingesting large portions sustained high response rates only at the beginning, supporting the idea that knowledge of an upcoming reward is not the same as its physiological ingestion (Ferriday & Brunstrom, 2011; Finlayson et al., 2007). Once participants consumed the accumulated snack and reached satiety, we observed a significant decrease in behavioral responding (Ferriday & Brunstrom, 2011; Pool et al., 2016; Rolls et al., 2007). Finally, we did not observe the strong habit-like behavior that sometimes arises in rodent tasks with delayed outcomes (Urcelay & Jonkman, 2019). Rather, participants remained goal-directed enough to cut back on instrumental responses once they recognized that the snack had lost its appeal (Tricomi et al., 2009).

In Experiment 2, we introduced a monetary reward condition—equally delivered in real time— to explore whether satiety cues or metabolic feedback might be critical in sustaining motivation. Although monetary stimuli initially invigorated instrumental behavior, they lost their motivational impact more quickly than food did, suggesting that even small, repeated food consumption can provide subtle physiological or sensory reinforcement that is absent in intangible rewards (Balleine & O’Doherty, 2010; Stice et al., 2010). Some theories suggest that intangible rewards, such as monetary gains, may remain stable over time because they are not subject to immediate biological signals (Eder & Dignath, 2016; Hogarth & Chase, 2011; Sescousse et al., 2013). However, our data point to a real-time discounting effect when monetary gains are dispensed trial by trial: in the absence of metabolic feedback, the monetary outcome appears more vulnerable to rapid devaluation once its novelty diminishes (Kirby & Marakovic, 1996; Madden et al., 2003). Hence, even though monetary cues elicited an initial transfer effect, food-based cues exhibited greater persistence because of their direct engagement with biological reinforcement mechanisms. Moreover, the action of retrieving and ingesting food probably provides additional reinforcing properties, such as hedonic taste, physiological satiety signals, and motor engagement—that intangible rewards lack (Pool et al., 2016; Rolls et al., 2007; Stice et al., 2008).

Notably, although the participants in the present study earned real monetary rewards, they did not physically retrieve or use them during the experiment. This lack of tangible consumption likely contributed to the more rapid decline in their motivational impact compared with that of food rewards. Prior research has shown that the physical act of eating—encompassing sensory pleasure, motor engagement, and physiological satiety signals—is crucial for maintaining incentive value (Berridge, 1996; Pool et al., 2016; Rolls et al., 2007; Stice et al., 2010). Indeed, our findings emphasize that immediate consumption is essential for capturing the dynamic interplay between hunger, satiety, and food cue-driven motivation. This understanding is particularly relevant when addressing overeating in environments with many food-related cues (Swinburn et al., 2011; van den Akker et al., 2018).

Importantly, our experiments aimed to observe devaluation—the decline in responding when the food becomes less appealing—rather than total extinction, where rewards are no longer available (Balleine & Dickinson, 1998; Dickinson, 1995). In this sense, although an extinction procedure was applied, the results suggest that the decline in cue-driven behavior was primarily facilitated by within-session satiety, which likely accelerated outcome devaluation. However, in line with our findings from Experiment 1, this process did not fully eliminate the behavior— indicating that satiety modulates but does not abolish cue reactivity. Nonetheless, cue-driven motivation persisted, revealing how robust Pavlovian associations can be (Corbit & Balleine, 2005; Watson et al., 2014). This finding supports the broader literature showing that while satiety may weaken responding, it rarely extinguishes cue‒outcome associations entirely (Holland, 2004; Panayi & Killcross, 2022). Future studies with longer or repeated training might crucially address whether short-term devaluation yields habit-like patterns resistant to satiety, as observed in some multisession experiments with rodents (De Wit & Dickinson, 2009; Hogarth & Chase, 2011; Neal et al., 2006). Examining these processes in individuals with obesity, who may exhibit altered cue reactivity, could refine our understanding of how these cues are devaluated in those populations (Meemken & Horstmann, 2019; van den Akker et al., 2018).

## Conclusions

In the present study, we found that satiety accelerated cue-driven devaluation in normoweight general population. Our experiments not only corroborate classic findings in animal models (Estes, 1943; Corbit & Balleine, 2005) but also extend recent evidence from human studies (Badioli et al., 2024; Cartoni et al., 2016; Colagiuri & Lovibond, 2015), showing that environmental cues retain motivational power despite outcome devaluation. These findings underscore the importance of incorporating physiological feedback in human paradigms involving food-cued learning to capture dynamic shifts in incentive value. The results suggest that large, immediate portions rapidly triggered satiety, corresponding to a sharp decline in responding, whereas smaller or delayed consumption prolonged approach behavior. However, in all cases, the participants eventually showed diminished motivation once satiated. Furthermore, we observed that although monetary outcomes also support PIT, they are devalued faster, perhaps due to the lack of involvement of biological reinforcements. Importantly, these results indicate that real-time ingestion enables a more precise understanding of how hunger transitions to satiety and thus modulates the persisting power of food cues.

## Supplementary material

Three separate bar-plots (mean ± SEM) summarize the raw data from all experimental conditions in both Experiment 1 and Experiment 2, for each of the three tasks. For the Pavlovian (Pedal block, Figure S1) and transfer tasks (Figure S3), the plots display (1) mean response rates, quantified as the number of key presses, and (2) mean latencies to the first key press, each averaged across all conditions and experiments. The third plot (Figure S2) depicts the mean response rates (number of key presses) on correct trials in the instrumental task, averaged by conditions across all five blocks for both Experiment 1 and Experiment 2.

**Figure S1.**
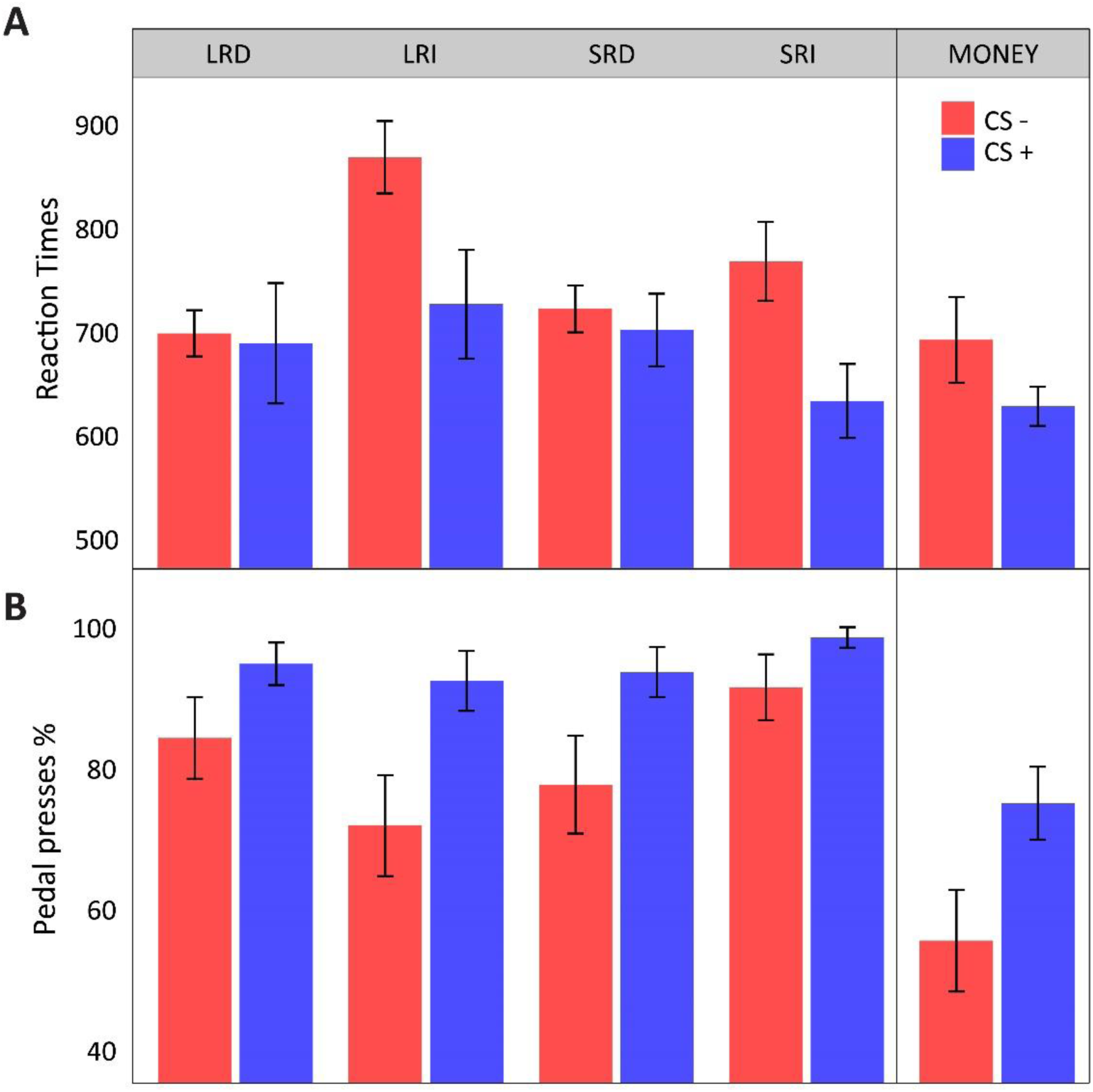
Pavlovian (Pedal block) task. Bar–plot illustrating mean latencies to the first key press (**A**), and mean response rates (number of key presses; **B**) ± SEM for each experimental condition in Experiment 1 and Experiment 2.

**Figure S2.**
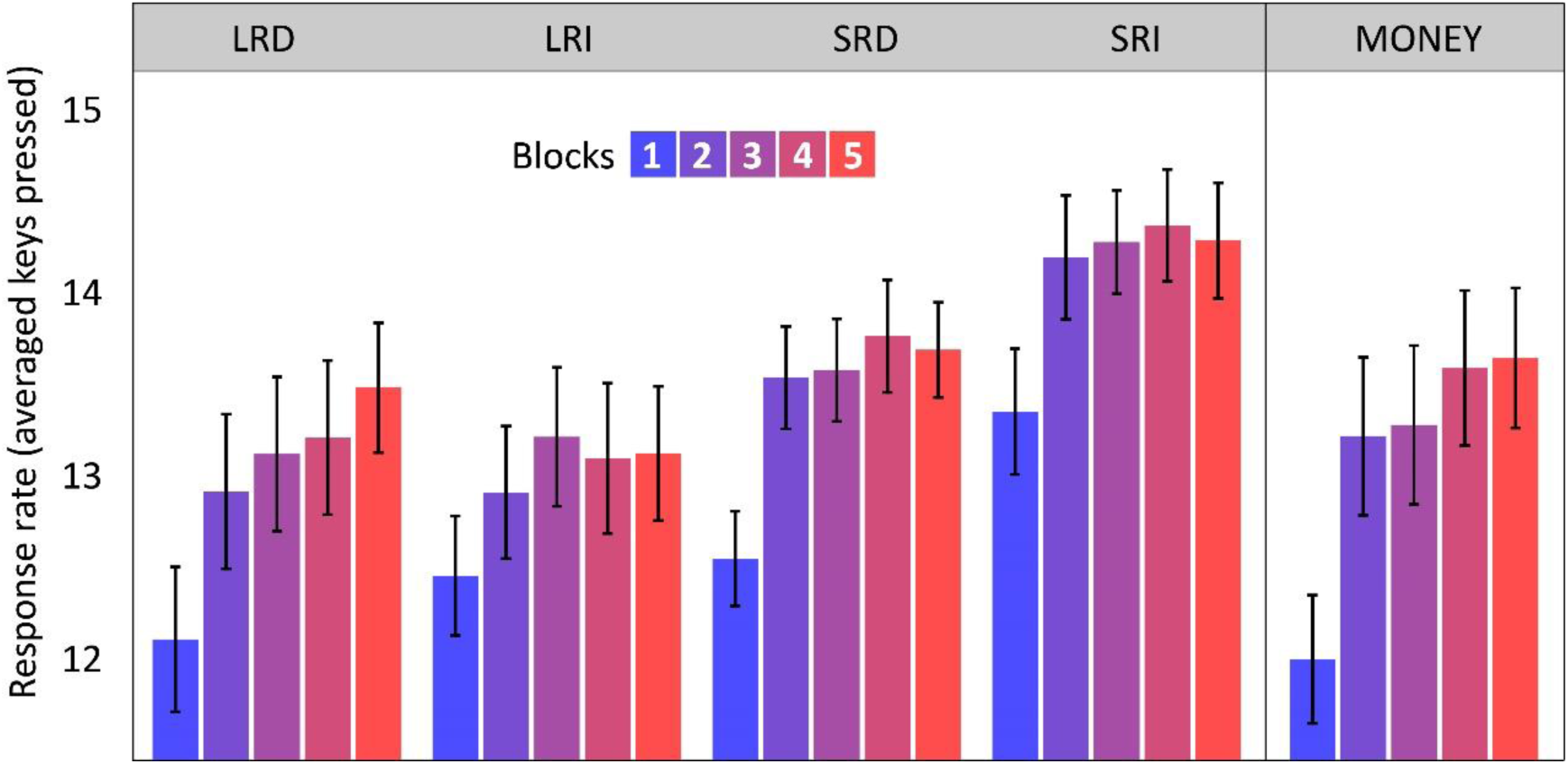
Instrumental task. The bar–plot shows mean response rates (number of key presses on correct trials) ± SEM across all five blocks of the task for each experimental condition in Experiment 1 and Experiment 2.

**Figure S3.**
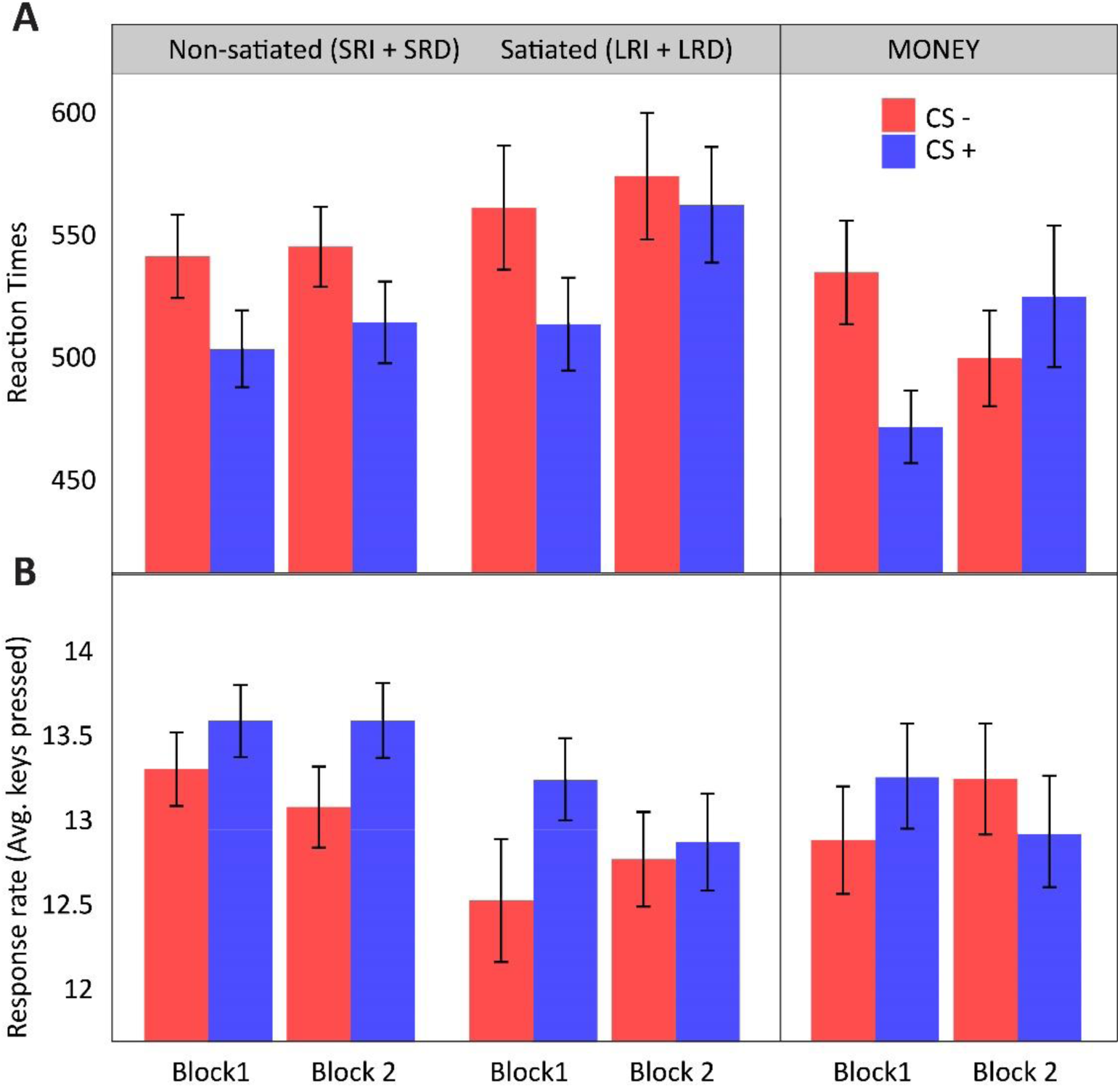
Transfer task. Bar–plot illustrating mean latencies to the first key press (ms; **A**), and mean response rates (number of key presses; **B**) ± SEM for each experimental condition in Experiment 1 and Experiment 2.

